# Influence of Wobbling Tryptophan and Mutations on PET Degradation Explored by QM/MM Free Energy Calculations

**DOI:** 10.1101/2024.04.30.591886

**Authors:** Anna Jäckering, Marc van der Kamp, Birgit Strodel, Kirill Zinovjev

## Abstract

Plastic-degrading enzymes, particularly poly(ethylene terephthalate) (PET) hydrolases, have garnered significant attention in recent years as potential eco-friendly solutions for recycling plastic waste. However, understanding of their PET-degrading activity and influencing factors remains incomplete, impeding the development of uniform approaches for enhancing PET hydrolases for industrial applications. A key aspect of PET hydrolase engineering is optimizing the PET-hydrolysis reaction by lowering the associated free energy barrier. However, inconsistent findings have complicated these efforts. Therefore, our goal is to elucidate various aspects of enzymatic PET degradation by means of quantum mechanics / molecular mechanics (QM/MM) reaction simulations and analysis, focusing on the initial reaction step, acylation, in two thermophilic PET hydrolases: LCC and PES-H1, along with their highly active variants, LCC^ICCG^ and PES-H1^FY^. Our findings highlight the impact of semi-empirical QM methods on proton transfer energies, affecting the distinction between a two-step reaction involving a metastable tetrahedral intermediate and a one-step reaction. Moreover, we uncovered a concerted conformational change involving the orientation of the PET benzene ring, altering its interaction with the side-chain of the ‘wobbling’ tryptophan from T-stacking to parallel π-π interactions, a phenomenon overlooked in prior research. Our study thus enhances the understanding of the acylation mechanism of PET hydrolases, in particular by characterizing it for the first time for the promising PES-H1^FY^ using QM/MM simulations. It also provides insights into selecting a suitable QM method and a reaction coordinate, valuable for future studies on PET degradation processes.

**TOC Graphic:** 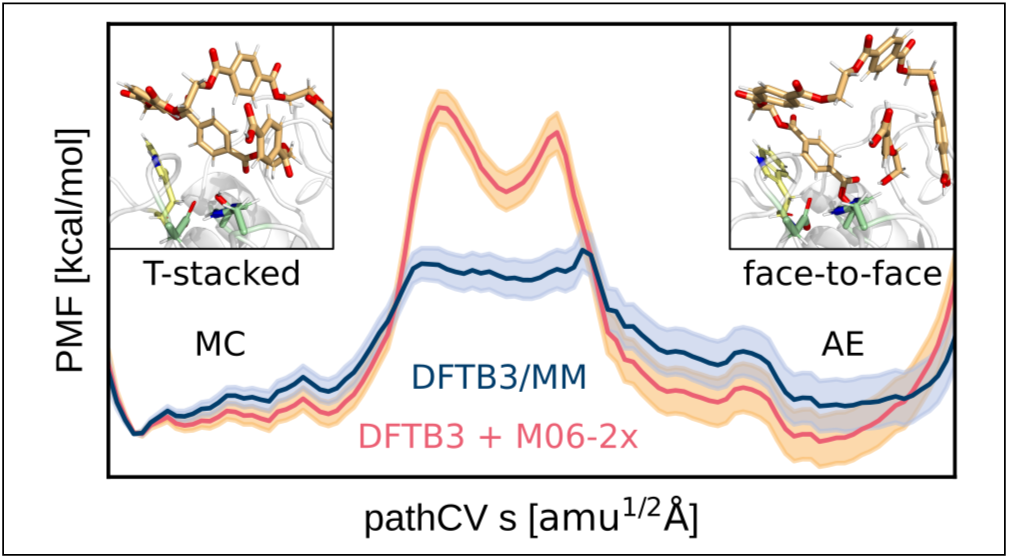

## 1 Introduction

Synthetic polymers, commonly referred to as plastics, have gained significant attraction in the global market due to their durability, affordability, and versatility.^1,2^ Presently, degradation methods for the immense amount of accumulated plastic waste are both environmentally and economically costly, pressing the need for an eco-friendly and efficient alternative. ^3^ Enzymatic degradation emerges as a promising solution, particularly focusing on poly(ethylene terephthalate) (PET), a widely used plastic in packaging and textiles. ^4,5^ The discovery of enzymes capable of PET degradation in 2005^6^ and the identification of *Ideonella sakaiensis* as an organism metabolizing PET^7^ in 2016 have paved the way for enzymatic degradation. Enzymes with PET-degradation function are often found among hydrolases (EC 3.1.1.x) and specifically cutinases (EC 3.1.1.74), but are now often categorized into the novel class of polyester hydrolases (EC 3.1.1.101).^7–9^ These enzymes share an α/β-hydrolase fold characterized by a core comprising eight β-strands and six α-helices with a surface-exposed catalytic triad typically composed of serine, aspartate, and histidine, facilitating PET hydrolysis. ^10,11^

Despite some wild-type (WT) enzymes exhibiting high activities, enzyme engineering is necessary to enhance efficiency and stability to facilitate industrial application, yielding two of the currently most active PET-degrading enzymes, the leave-branch compost cutinase F243I/D238C/S283C/Y127G (LCC^ICCG^) and polyester hydrolase I L92F/Q94Y (PES-H1^FY^) variants.^5,12,13^ Notable, these two variants share a mutation at the same position, yet with divergent effects: activity of LCC could be enhanced by elimination of tyrosine (Y127G) in addition to other mutations present in the LCC^ICCG^ variant, while an introduction of tyrosine (Q94Y) in combination with introduction of phenylalanine in close proximity yielded the PES-H1^FY^ variant with increased activity.^5,13^ We previously found that the entry of PET into the active binding cleft and the subsequent Michaelis complex (MC) formation is promoted by the LCC F243I/Y127G (IG) variant^14^ but the precise effect remains unclear and demands elucidation of the energy profile of the first reaction step, the acylation, to fully understand MC stabilization.

The catalytic hydrolysis reaction of several PET-degrading enzymes was investigated using quantum mechanics/molecular mechanics (QM/MM) simulation studies, which enable investigation of the region contributing to the target reaction at the QM level for bond breaking and formation, while managing the computational demand by treating the remaining part of the system at the MM level.^15–17^ These studies revealed two predominant binding poses of PET: pro-*S* ^18–20^ and pro-*R*.^5,21^ An energetic preference for catalysis starting from the pro-*S* pose is proposed, which allows for T-stacking π-π interactions between PET’s benzene ring and a conserved tryptophan positioned at one end of the catalytic binding cleft, which is also called ‘wobbling’ tryptophan (LCC: W190, PES-H1: W155). ^15,22^ The catalysis mechanism after substrate binding can be divided into two main processes: acylation and deacylation (Fig. 1).^15–17,23–26^

**Figure 1:**
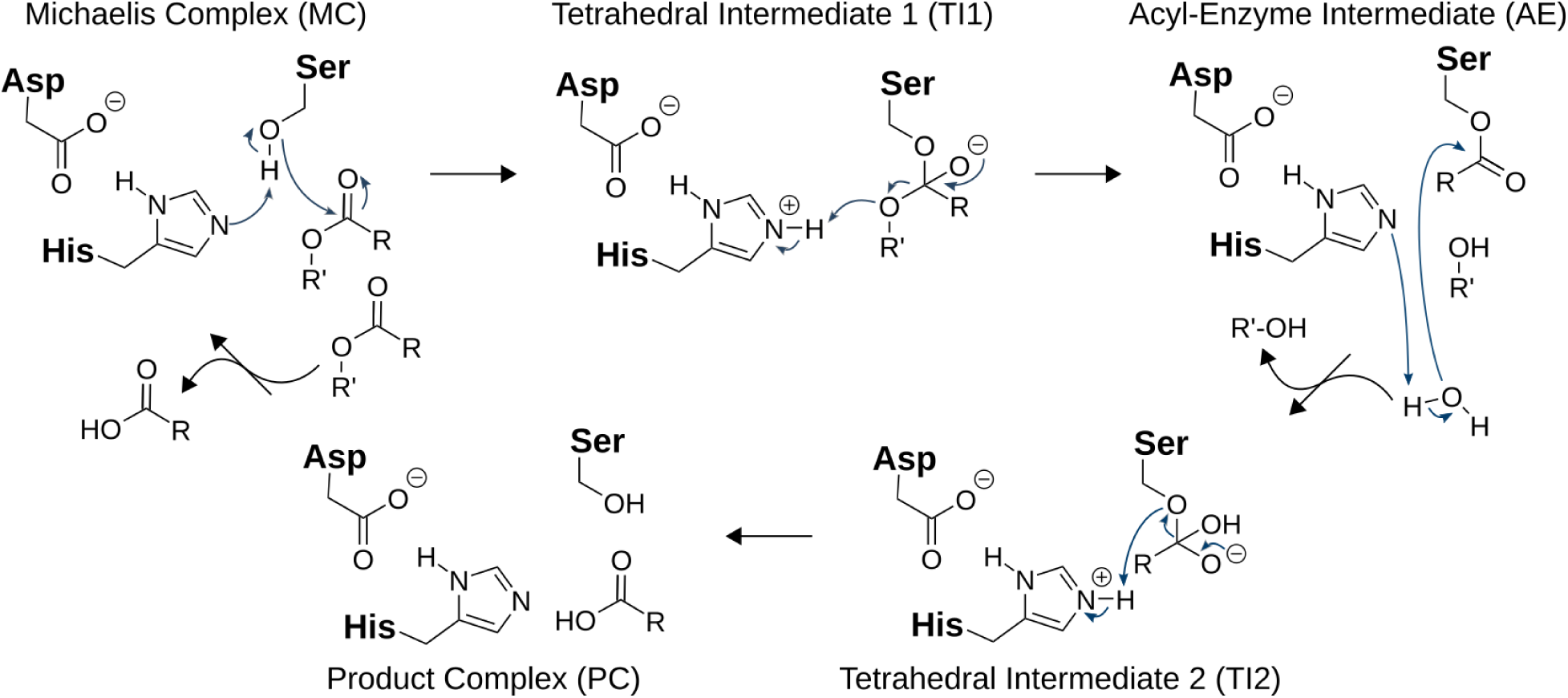
Mechanism of PET-degradation as proposed by Boneta *et al.*^15^ The mechanism can be divided into two steps: First, the acylation of the Michaelis complex (MC) yielding the acyl-enzyme intermediate (AE) and second, the hydrolysis yielding the product complex (PC), both incorporating a tetrahedral intermediate (TI).

During acylation, the catalytic serine’s nucleophilicity increases via proton transfer to the catalytic histidine, which is polarized by the catalytic aspartate. This facilitates nucleophilic attack on the ester’s carbonyl carbon, with the resulting oxyanion stabilized by the oxyanion hole. Subsequent cleavage of the ester bond yields the acyl-enzyme intermediate (AE), releasing an alcohol molecule from the active site after a proton transfer from the catalytic histidine to the resulting oxyanion. In deacylation, a water molecule’s oxygen nucleophilically attacks the carbonyl carbon of AE, generating a second tetrahedral intermediate (TI) and reprotonating the catalytic histidine. Finally, the AE bond breaks, yielding the second product with a carboxyl group, while the serine is reprotonated, restoring the initial state. Products comprise smaller PET oligomers, as well as mono(2-hydroxyethyl) terephthalic acid (MHET), bis(2-hydroxyethyl) terephthalic acid (BHET) and the final monomers terephthalic acid (TPA) and ethylene glycol (EG).^27–29^

There is debate over whether acylation and deacylation occur in one or two steps, with the TI potentially representing a tetrahedral transition state (TS).^15,16^ Identifying the rate-determining step remains challenging, with some studies suggesting acylation and others deacylation.^15,17,23–26,30,31^ Product diffusion from the active site, necessary for subsequent substrate hydrolysis, may also be rate-limiting, as suggested by Shrimpton-Phoenix *et al.*,^31^ though other simulations contest this notion. ^26^ Various studies discuss the catalytic histidine’s polarisation mechanisms, considering proton transfer in addition to pure polarisation.^20,25,31^ QM/MM studies highlight the role of specific amino acids (Ser, Ile, Ala) near the catalytic aspartate in stabilising the histidine, influencing proton transfer dynamics.^23,32^ The choice of the QM region significantly impacts observations, with larger regions offering greater accuracy but escalating computational costs. Mutation experiments confirm the importance of isoleucine, with alanine substitutions resulting in activity loss.^23^ Such insights are crucial for evaluating mutation sites, particularly in enzyme redesign aiming to lower the free activation energy by stabilising transition states, beyond substrate binding. ^33,34^

Another interesting residue is a conserved tryptophan, the ‘wobbling’ tryptophan, which exhibits three different conformations in the mesophilic *Ideonella sakaiensis* PETase (*Is*PETase), while maintaining a single conformation in other, mostly thermophilic polyester hydrolases.^20,35^ This rigidity is attributed to steric hindrance by a nearby histidine, which in *Is*PETase is substituted by a serine. Extensive MD and metadynamics simulations, coupled with experimental analyses, have revealed that the presence of this serine and an adjacent isoleucine in *Is*PETase, instead of histidine and phenylalanine in other PET-degrading enzymes, enhances the flexibility of the tryptophan and loop regions within the active site, positively impacting activity.^36,37^ Moreover, the conformational flexibility of tryptophan allows T-shaped π-π interactions with PET’s benzene ring, thereby expanding the space within the active site and favoring PET binding. Following hydrolysis, suggested conformational changes of the PET benzene ring lead to energetically less favored face-to-face stacking and promote product release.^20,35,37^

The detailed impact of the ‘wobbling’ tryptophan on the reaction energetics remains unexplored. Moreover, prior publications propose inconsistent conclusions regarding whether the acylation and deacylation mechanism occurs as a one- or two-step process. ^15–17,26^ To address these questions, we aim to elucidate the conformational changes of the benzene ring of PET that alter its interaction with the ‘wobbling’ tryptophan, and to shed light free energy profile of the acylation reaction to distinguish between one- and two-step mechanisms.

For this reason, we performed QM/MM simulations of the acylation step of PES-H1 and LCC as well as their variants, PES-H1^FY^ and LCC^IG^, the latter representing LCC^ICCG^ excluding the stabilizing disulfide bridge. This is the first QM/MM investigation of PES-H1, and provide additional insights into the LCC, which has been analyzed in several previous studies and thus allows for comparison and validation of our results. ^15,17,22,24,32^ The QM/MM method is used in combination with the adaptive string method (ASM),^38^ which offers a flexible approach for incorporating collective variables to delineate the reaction pathway. This methodology sheds light on the previously overlooked impact of the ‘wobbling’ tryptophan, addressing gaps in our understanding of PET degradation mechanisms, thus helping to drive the development of industrially applicable enzymes.

## 2 Methods

### 2.1 Parameterisation

The parameterisation of a small PET analogue, comprised of three units, has been previously documented by Pfaff and colleagues. ^13^ To model the analogue of AE, three residues additional were parameterised using the same methodology applied to PET. These residues specifically represent the serine bonded to PET and two residues corresponding to the units subsequent to the cleavage of the connecting ester bond.

The parameterisation procedure involved quantum mechanics calculations conducted at the HF/6-31G* level using Gaussian 09.^39^ This was undertaken to establish generalised AMBER force field (GAFF)^40^ parameters for both the terminal and central units. Subsequently, restrained electrostatic potential (RESP) calculations^41,42^ were executed through the antechamber tool^43,44^ available in AmberTools 21.^45^ This step aimed to derive partial charges, which were then redistributed using prepgen to achieve a zero net charge for each PET unit. Finally, input files for the GROMACS simulation software^46^ were generated using the ACPYPE tool.^47^

### 2.2 Preparation of Input Structures

For each enzyme, namely LCC and PES-H1, the pro-*S* conformation with the minimal distance between the PET ester carbon and the catalytic serine’s *γ*-oxygen was extracted from the WT HREMD simulations, as detailed in a previous publication. ^14^ These starting structures comprised a single PET molecule involving five units besides the enzyme. To ensure compatibility with the AMBER software utilised in this study, the system was resolvated and setup using *tleap*, as the solvated box generated from GROMACS^46^ is not congruent with the box types employed in AMBER.^48^ Starting structures for the variants were obtained by introducing mutations to the WT structures, ensuring similar PET binding poses for improved comparability. The system was solvated and neutralised by the addition of TIP3P water and sodium or chloride ions, respectively. The system was represented using the AMBER14SB force field, and parameters for all three PET unit types were incorporated using *tleap* as implemented in AMBER. The protonation states of titratable amino acids were maintained as originally suggested by Propka 3.4, ^49,50^ resulting in a net charge of +5 for LCC and LCC IG, and –5 for PES-H1 and PES-H1 FY.^14^ The final system’s energy was locally minimised using the steepest descent algorithm^51^ followed by the conjugate gradient algorithm.^52^ Subsequent equilibration runs were performed at a temperature of 303 K and pressure of 1 bar in the NVT and NPT ensemble, utilising the Langevin thermostat^53^ and the Berendsen barostat,^54^ respectively, with restraints applied to the position of the protein and PET. The production run spanned 20 ns in the NPT ensemble, incorporating one-sided harmonic distance restraints to maintain the PET ester in a productive pose. The harmonic restraint, up to 20 kcal/mol/Å^2^, was initiated for a distance above 3.5 Å between the catalytic serine’s *γ*-oxygen and PET’s ester carbon, and for a distance above 3 Å between PET’s ester carbonyl-oxygen and the centre of mass of the backbone amine’s nitrogen of the oxyanion hole residues, respectively. This restraint becomes linear for distances above 10 Å. A periodic box was employed for all runs, SHAKE algorithm was used^55^ to constrain hydrogen bonds. To reduce computational demand, a cutoff at 8 Å for short-range non-bonded interactions was introduced, and the PME method^56,57^ was applied to treat electrostatic interactions.

The AE was created from the reactant’s structure after the MM production run using PyMOL, retaining water molecules, ions, and box dimensions. The same protocol as for the reactants was followed to obtain the preferred PET benzene ring orientation and corresponding tryptophan–PET dihedral angle from the 20 ns production run, in which the CV distances were restrained to keep the two PET units adjacent to the broken ester bonds within the AE basin. As the overall conformation of PET was altered due to conformational changes of the remaining units of the long, flexible PET chain during the MM production run, the AE structure was subjected to the QM/MM preparation right after MM minimisation diminishing the MM equilibration and MM production run steps. This involved a QM/MM optimisation using steepest descent and conjugate gradient for both, reactant and AE. Only for the AE, a 50 ps QM/MM run was performed in the NVT ensemble applying a one-sided harmonic restraint to adjust the tryptophan–PET angle to the prevalent one during the MM production run. For the PES-H1 FY variant, this step was skipped as the preferred tryptophan–PET angle was already obtained after MM MD and was, thus, already existent after QM/MM optimisation. The last step before the actual ASM run comprised an unrestrained 50 ps QM/MM relaxation in the NVT ensemble reactant and AE. The QM region comprised 66 atoms, including the two PET units connected by the scissile ester bond, and the sidechains of the catalytic serine, histidine, and aspartate (Fig. 2). It was described by the DFTB3 method without using the SHAKE algorithm to allow for proton transitions.^58^ The non-bonded cutoff was increased to 10 Å for minimisation and to 9,999 Å for relaxation.

**Figure 2:**
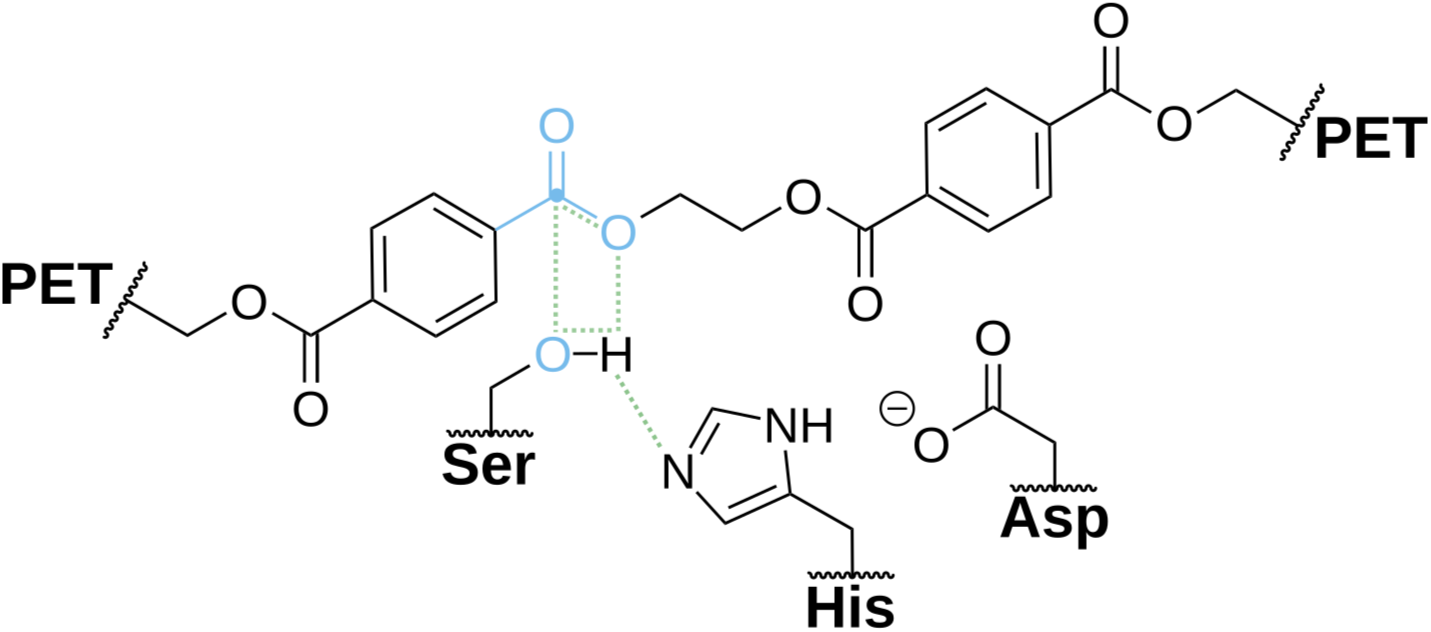
QM region and CVs employed in this study. The QM region includes two PET units connected via the scissile ester bond and the side chains of the three catalytic triad residues (PES-H1: S130, D176, H208; LCC: S165, D210, H242). The distance CVs are highlighted in green, and the point-to-plane CVs are in blue, with the PET ester’s carbon corresponding to the point marked by a blue circle. The remaining atoms forming the plane include those in the ester between the PET units in the MC and the ester between PET and the catalytic serine in the AE, respectively. An additional dihedral CV is included, which is highlighted in figure 4.

### 2.3 ASM

The analysis of PET deacylation was conducted using the ASM, as implemented in AMBER.^38^ The initial and final states for the pathway connecting the reactant and the AE in CV space were defined through QM/MM relaxation simulations. Five relevant distances describing the reaction and one dihedral representing the ring orientation change were employed as CVs, as well as two point-to-plane distances reflecting the hybridisation state of the PET ester’s carbon to enhance convergence (Fig. 2). MD simulations were initiated from a total of 96 nodes, equidistantly positioned along the initial guess. Half of these simulations started from the reactant state, while the other half began from the AE state. The structures were brought to the corresponding node positions during 1 ps by gradually increasing the force constants from zero to the target value.

Upon convergence of the string to the MFEP, a path-CV was defined, illustrating the progress along the MFEP and consequently, the advancement along the reaction. umbrella sampling MD (USMD) simulations^59,60^ were then performed, restraining each replica to the corresponding node on the string for the US windows using an automatically defined potential. The PMF was obtained via umbrella integration (UI),^61–63^ and assumed to be converged when the 95% confidence interval at the TS fell below 1 kcal*·*mol^-1^.

### 2.4 DFT correction

The PMF obtained from the ASM runs was corrected to M06-2x/6-31+G(d,p) level of theory^64^ by applying a cubic splines correction based on single point energy estimates for relaxed structures along the reaction path. Single-point calculations were carried out for the resulting optimised structures using the DFTB3 method and, additionally, the M06-2X functional^64^ with the 6-31+G(d,p) basis set, employing the Gaussian 09 software. ^39^ The energy difference (Δ*E* = *E_DFT_ − E_DFTB3_*) was then interpolated and subtracted from the original PMF values to obtain the corrected PMF.

## 3 Results and Discussion

### RC and AE prefer different PET ring orientations

To conduct QM/MM simulations of PET acylation, we extracted starting structures of PET–enzyme complexes from Hamiltonian replica exchange MD (HREMD) simulations as detailed in our previous work.^14^ These structures involved a PET chain with 5 units and either LCC or PES-H1. The initial configurations for the LCC^IG^ and PES-H1^FY^ variants were derived by introducing corresponding mutations to the WT enzymes, ensuring a consistent starting point for comparison. Analysis of the binding poses revealed that the PET units sharing the scissile ester bond are tightly bound, whereas units further away exhibit looser interactions with the enzyme. This contradicts assumptions of simultaneous binding of several PET units.^65^

Four resultant Michaelis complexes (MC)s underwent 20 ns MM simulations, confirming the stability of chosen binding poses. However, chain termini remained flexible (Fig. S1). The final structures served as templates to manually generate AE structures using PyMOL. Subsequently, they were subjected to 20 ns MM simulations with restraints to prevent the leaving group from diffusing away from the AE. Two notable findings emerged: Firstly, loosely bound PET units exhibited greater flexibility compared to those near the scissile ester bond, resulting in conformations unsuitable for subsequent QM/MM simulations using the ASM due to the difference to MC conformations (Fig. 3).

**Figure 3:**
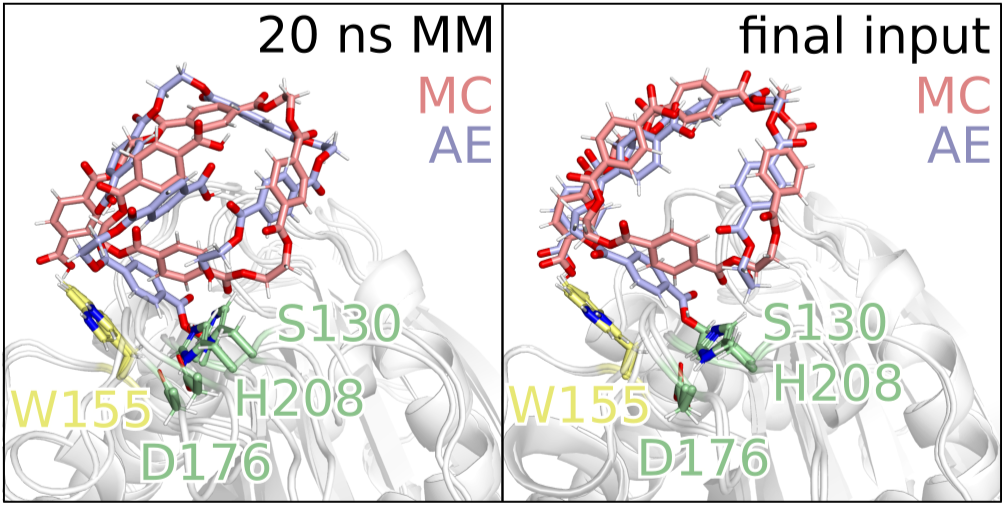
PET conformations of the MC (red) and AE (blue) for PES-H1 after 20 ns MM production run (left) with prior two-step equilibration and the final structures used as input for the ASM run (right). Here, an additional QM/MM optimisation and 50 ps relaxation followed the 20 ns production run for the MC. The AE was generated from the MC structure after 20 ns MM production run and subjected to MM minimisation, QM/MM optimisation and two 50 ps QM/MM simulations, of which the first included restraints to set the dihedral angle to that observed during the 20 ns MM production run while the second was performed without restraints to relax the structures prior to ASM.

Secondly, the benzene ring of PET bound to the catalytic serine in the AE demonstrated a preference for a parallel orientation with respect to the ‘wobbling’ tryptophan, whereas the MC favoured a T-stacked orientation. This transition from T-stacked to parallel π-π interactions was observed during MM minimisation (PES-H1^FY^) and during the beginning (LCC, LCC^IG^) or middle (PES-H1) of the MM production run. This shift, previously discussed by Han *et al.*^20^ in the context of product liberation, was not specifically mapped to a reaction step. Our results suggest that this transition occurs during the acylation phase. To address these findings, we conducted MM minimisation for AE structures without equilibration or 20 ns production runs prior to QM/MM minimisation. This was followed by an additional 50 ps QM/MM run with restraints to enforce the transition of the benzene ring from T-stacked to parallel. However, using plane angles as restraints or CVs in the ASM run is currently not implemented. We therefore identified a dihedral angle that correlated well with the plane angle, serving both as a restraint in structure preparation and as a CV during ASM reaction simulation (Fig. 4). An angle of approximately 0° corresponds to a T-stacked orientation, while an angle of approximately 90° indicates a parallel orientation. After enforcing the preferred parallel plane–plane angle of the AE, both MC and AE structures, were subjected to a 50 ps QM/MM relaxation without restraints before conducting the final ASM run.

**Figure 4:**
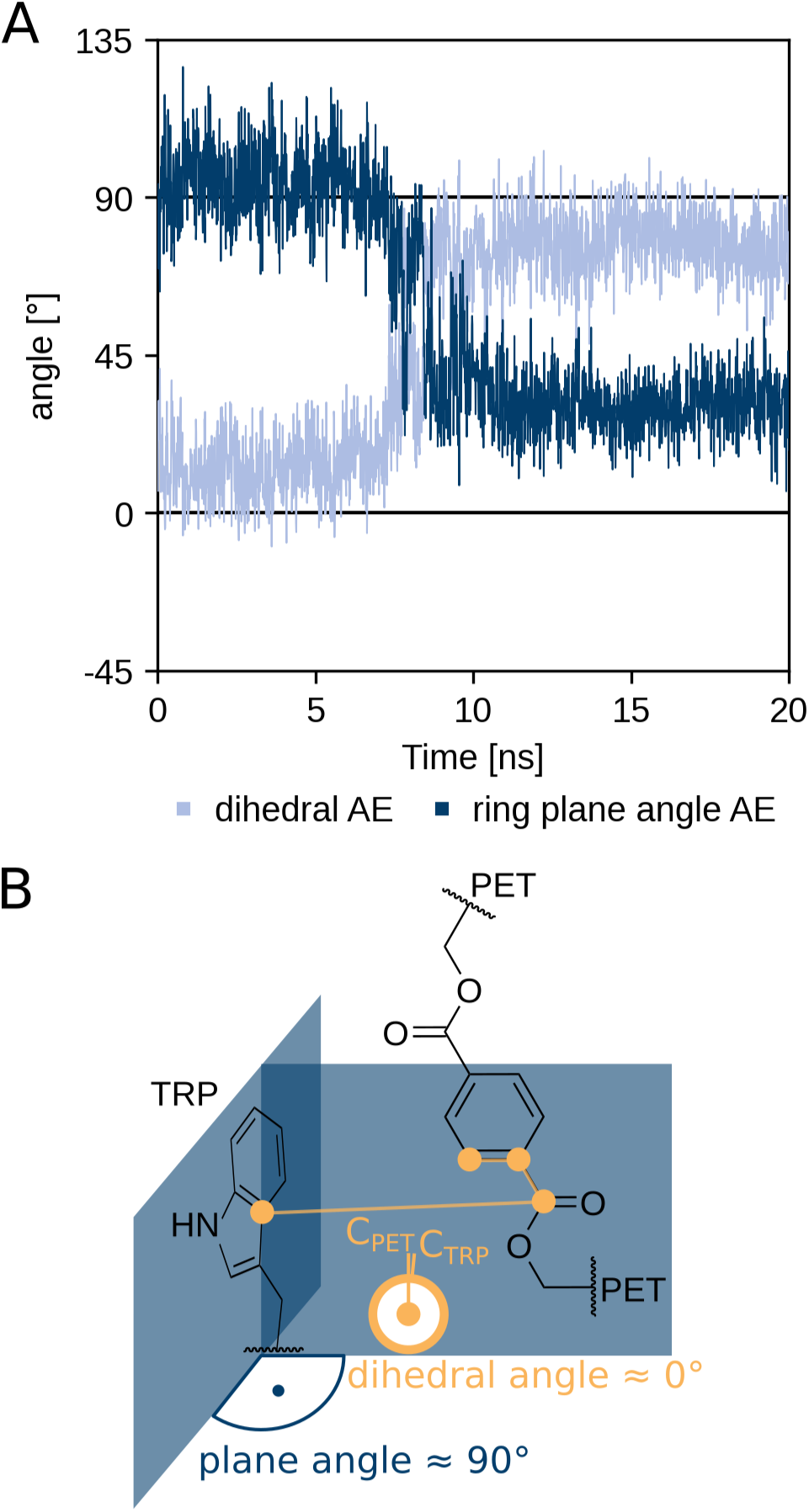
Dihedral angle and the angle between the tryptophan and PET ring planes during the 20 ns MM production run of the AE (lightblue, darkblue) of PES-H1 (A). The definition of the dihedral (orange) and the ring plane angles (blue) is highlighted schematically for the MC state preferring an approx. T-stacked orientation with a plane angle of approx. 90°, which correlates with a dihedral angle of approx. 0° (B).

### PET ring orientation change correlates with acylation

To perform QM/MM calculations, we used the final MC and AE structures after 50 ps QM/MM relaxation as input configurations (Fig. S2). We employed the semi-empirical DFTB3 method^58^ to describe the QM region, encompassing the side chains of catalytic residues (LCC and LCC^IG^: S165, D210, H242; PES-H1 and PES-H1^FY^: S130, D176, H208) and the two PET units sharing the scissile ester (Fig. 2). This QM method, successfully applied in previous studies in combination with ASM, offers a balanced trade-off between accuracy and computational demand. ^16^ Additionally, we corrected the PMF on DFT level using the M06-2X functional^64^ with the 6-31+G(d,p) basis set, which has proven effective in QM/MM studies of PET hydrolases. ^15,17^ We used relevant distances depicting bond formations and breaking during the reaction as CVs, alongside two point-plane distances defining the hybridisation state of the PET ester’s carbon, to enhance convergence.^16^ Furthermore, we included the dihedral defining the plane angle change as a CV. Average values for each CV, as well as the plane angle were calculated for each node after the ASM string was converged (Fig. 5 and 6).

**Figure 5:**
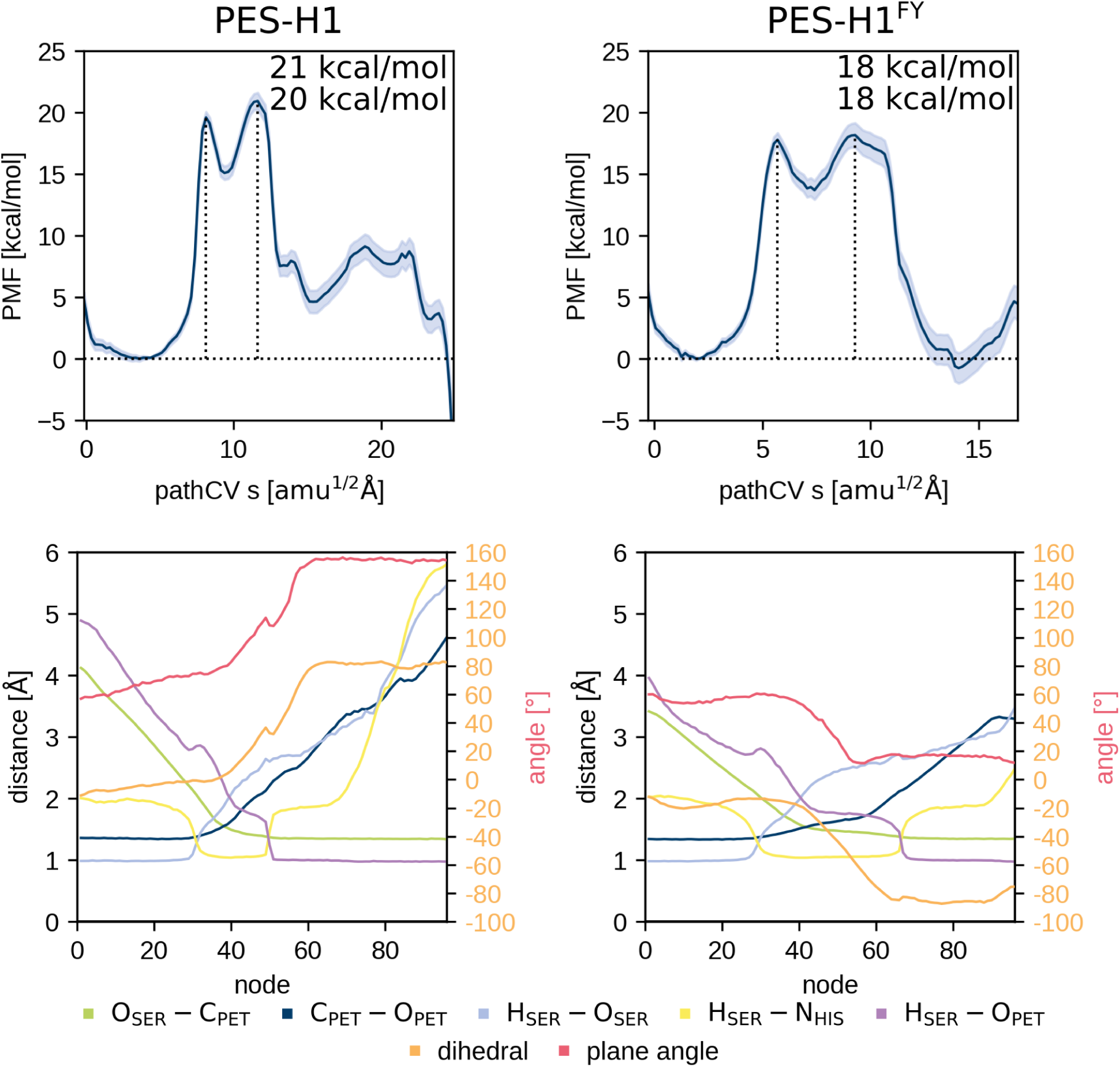
PMF of the PET deacylation of PES-H1 (left) and its highly active variant PES-H1^FY^ (right). The height of the two barriers is provided as well. The lower plots show the corresponding averaged distance CV values per node during umbrella sampling of the converged string and, thus, depicts the CV progression along the reaction. Node 0 corresponds to the MC and node 96 to the AE state. Additionally, the angle between the ring planes of the ‘wobbling’ tryptophan and the PET benzene moiety is plotted in red to highlight the correlation with the dihedral angle change in orange.

**Figure 6:**
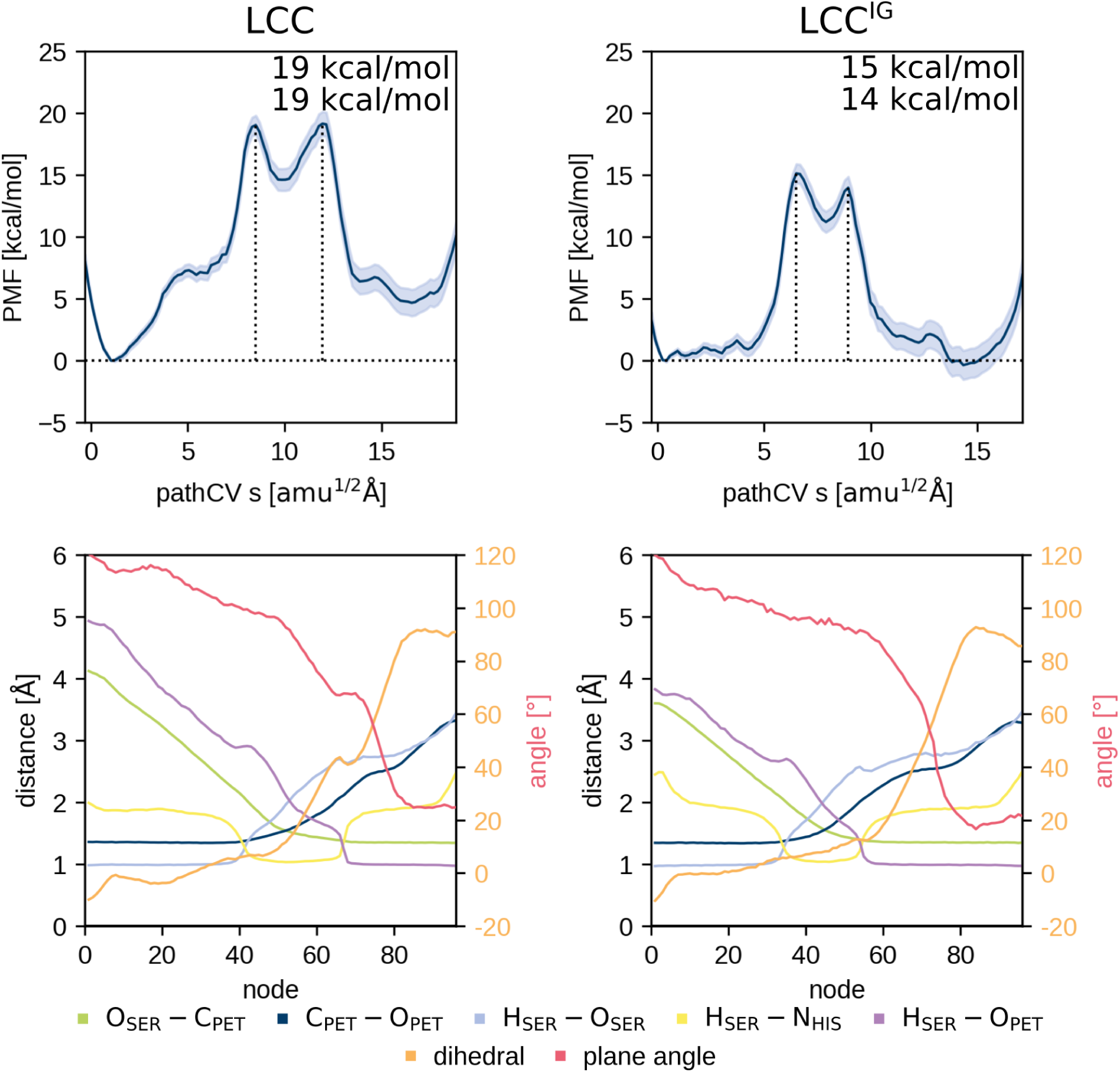
PMF of the PET deacylation of LCC (left) and its highly active variant LCC^IG^ (right). The height of the two barriers is provided as well. The lower plots shows the corresponding averaged distance CV values per node during umbrella sampling of the converged string and, thus, depicts the CV progression along the reaction. Node 0 corresponds to the MC and node 96 to the AE state. Additionally, the angle between the ring planes of the ‘wobbling’ tryptophan and the PET benzene moiety is plotted in red to highlight the correlation with the dihedral angle change in orange.

A smooth transition from approximately 0° (T-stacking) to approximately *±*90° (parallel orientation) towards the ring plane of the ‘wobbling’ tryptophan was revealed, showing good correlation with the plane angle change during the reaction. This smooth transition underscores the importance of including the plane angle change as a dihedral CV. As a proof of concept, we performed the same ASM calculation for PES-H1^FY^ without including the dihedral CV. Here, the dihedral for MC and AE was not fixed, allowing for more flexibility and a mixture of parallel and T-stacked conformations when approaching the product. This T-stacked conformation is adopted after 180° rotation, so that the PET ring atom defining the dihedral points away from the ‘wobbling’ tryptophan instead of towards it as in the MC (Fig. 7). Thus, the change in ring orientation happens even when it is not included in the CVs, highlighting the link between this change and the reaction.

**Figure 7:**
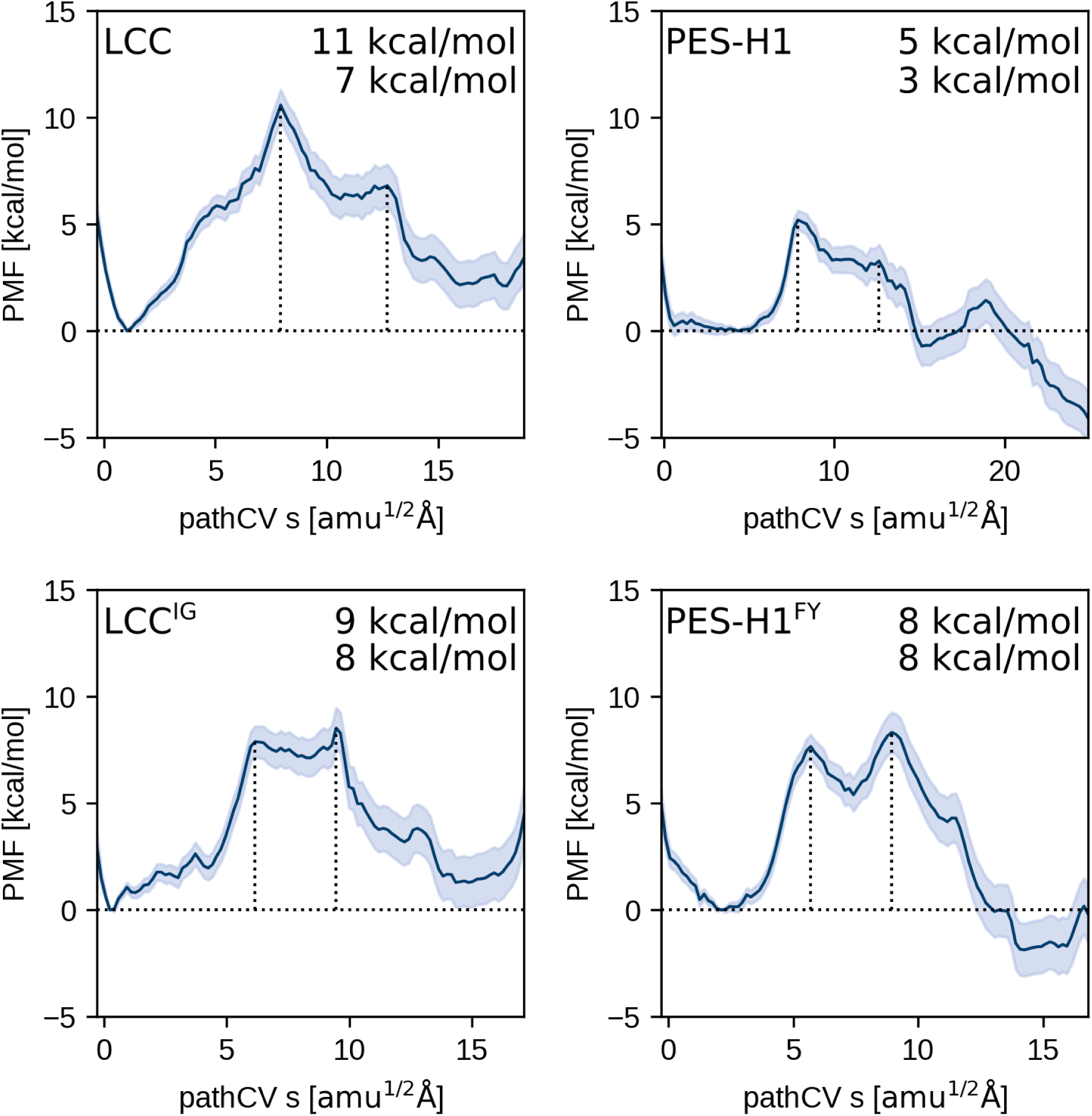
Comparison of the PMF using DFTB3 only (top) and after correction for DFT level (center), as well as the average dihedral and plane angles per node (bottom) for PES-H1^FY^ when incorporating the dihedral as a CV (A) and without the dihedral CV (B).

The progression of the distance CVs in combination with the PMFs describes the course of the reaction and enables mapping the ring orientation change to a specific step (Fig. 5 and 6). The reaction mechanism matches that from previous works, suggesting a two-step acylation with the tetrahedral intermediate state a shallow minimum on the free energy surface.^15,17,22–25,30,32^ Firstly, the catalytic histidine abstracts a proton from the catalytic serine, resulting in TS1. Then, the serine attacks the ester carbon, yielding the tetrahedral intermediate with its oxyanion stabilised by the oxyanion hole. The oxyanion is then converted back into a carbonyl oxygen by breaking the ester bond between the two PET units, generating a new oxyanion. The second transition state corresponds to the second proton transfer from the catalytic histidine to this oxyanion of the leaving group, liberating an alcohol and resulting in the AE. The steepest change of the plane and dihedral angles begins with the formation of the tetrahedral intermediate, and in the case of LCC^IG^, starts after the second proton transfer, and is thus concerted with the acylation (Fig. 5 and 6). Comparing the PMF of the ASM runs with and without the dihedral CV shows that the barriers in the corrected PMFs are somewhat larger when the dihedral CV is not included, suggesting that the conformational change during reaction is energetically favourable, which is neglected when the corresponding CV is omitted (Fig. 7).

### Over- or underestimation of proton transfers by different QM methods influences the energetics of the reaction

Comparison of the uncorrected DFTB3 PMF profiles with those corrected by DFT (M06-2X/6-31G(d,p)) shows that the energy barriers are generally much lower for DFTB3, most likely due to underestimation of the energy barrier for proton transfer (Figs. 7 and S3). In general, the choice of the QM method influences the PMF outcome, which is mirrored by the diverging results in previous works, suggesting either a one-step^16,26^ or a two-step^15,17,22–25,30–32^ mechanism for the acylation of PET hydrolases. As an example, Boneta *et al.* used the AM1^66^ method to describe the QM region and corrected the resulting PMF on DFT level using the M06-2X functional^64^ with the 6-31+G(d,p) basis set as in this study, which yielded a clear two-step PMF for LCC^ICCG^ and *Is*PETase, potentially as AM1 tends to overestimate energy barriers.^15^ In turn, García-Meseguer *et al.* obtained a one-step profile for the FAST-PETase using the DFTB3 method^58^ which tends to underestimate proton transfer barriers. ^16^ Similarly, Jerves *et al.* postulated a one-step profile using the Perdew–Burke–Ernzerhof (PBE) functional, ^67,68^ which also gives small energy barriers for proton transfers.^26,69,70^ The PMFs obtained here using the DFTB3 method also suggest that the reaction might be a one-step rather than a two-step mechanism (with either a very shallow or no significant minimum for the TI), but it is turned into a clear two-step profile upon correction to DFT (M06-2X) (Figs. 7, S3, 5 and 6) Thus, we suggest a two-step mechanism including a metastable intermediate state for the acylation stage of PET degradation, which may appear as a one-step mechanism when using methods underestimating proton transfer energies such as DFTB3 and PBE.

### Decreased acylation barrier as one reason of increased activity of variants LCC^IG^ and PES-H1^FY^

Our previous study analysed the effect of single residues on the entry of PET into the active site of LCC and PES-H1 and the corresponding high activity variants LCC^ICCG^ (represented by LCC^IG^) and PES-H1^FY^.^14^ The free energy surfaces suggested that the mutated residues promote PET entry by facilitating an unhindered entry and by stabilising the bound over the unbound state. Our results suggest that both variants exhibit significantly lower activation free energy barriers for acylation (considering a 95% confidence interval of *≈*1 kcal*·*mol^-1^; Fig. 5 and 6). Furthermore, the MC shows the same or higher energy as the AE in the corrected and uncorrected PMF of the variants, while it exhibits a lower energy in the WT profiles suggesting either a destabilisation of MC or a stabilisation of the AE upon mutation. A destabilisation of the MC would simultaneously explain the reduced activation free energy barrier with respect to the MC, while stabilisation of the AE can result from increased stabilisation of the TS. A stabilisation of the MC upon mutation was suggested for all mutation sites of both variants, LCC^IG^ and PES-H1^FY^, based on free energy surfaces in a previous study.^14^ Thus, we suggest that a stabilisation of the TS and AE (with respect to MC) is the cause of the changes in the PMF and the reduction in activation free energy barrier, promoting acylation and potentially also the PET-degradation activity of the variants as found experimentally.^5,13^

### 3.1 Conclusion

This study explored the initial chemical step of enzymatic PET degradation, the acylation process, in detail, using QM/MM MD sampling with the adaptive string method (ASM). We focused on two well-studied PET hydrolases, LCC and LCC^ICCG^ (abbreviated as LCC^IG^), alongside PES-H1 and PES-H1^FY^.^5,13^ Our investigation revealed two distinct conformations of the PET benzene ring, facilitating different types of π-π interactions with a ‘wobbling’ tryptophan residue in the MC (T-stacked) and AE (parallel). This conformational shift appears to occur concertedly with acylation and contributes to the resulting free energy profile.

Therefore, it is crucial to consider the varying orientations of the PET benzene ring during QM/MM simulations (e.g. by including a CV describing this during sampling), an aspect previously overlooked. Furthermore, we found that acylation proceeds through a two-step process involving a metastable tetrahedral intermediate, which is potentially missed in other studies due to the QM method employed. Additionally, both high-activity variants exhibited reduced activation free energy barriers compared to their respective WT enzymes. Interestingly, in the variants, the MC and AE states were energetically equivalent, whereas the MC is more stable in the WT enzymes. We hypothesise that this relative stabilisation of the AE is coupled with the stabilisation of the transition state, contributing to enhanced activity. However, we assume that the reduction of the energy barrier by approx. 2-5 kcal/mol is not the only factor driving the substantial activity increase. The multifaceted nature of PET enzyme activity is influenced by various mechanisms such as substrate adsorption, entrance to the binding site, productive substrate binding, and product liberation, in addition to the catalysis of the chemical process of substrate conversion addressed here. Consequently, comprehending the heightened PET-degradation activity of LCC^ICCG^ and PES-H1^FY^ demands a comprehensive approach encompassing all pertinent mechanisms.

## Acknowledgement

We gratefully acknowledge the computing time granted through the hybrid computer cluster purchased from funding by the Deutsche Forschungsgemeinschaft (DFG, German Research Foundation) project number INST 208/704-1 FUGG, and the Center for Information and Media Technology at Heinrich Heine University Düsseldorf as well as the computational facilities of the Advanced Computing Research Centre, University of Bristol. A.J. thanks the German Federal Environmental Foundation for the ideal and financial support. M.v.d.K and K.Z. thank the Engineering and Physical Sciences Research Council (grant EP/V011421/1). KZ acknowledges a Maria Zambrano fellowship by Ministerio de Universidades (Spain).

## Supporting Information Available

The following files are available free of charge.

- PET conformations after 20 ns MM production run
- PET conformations after 50 ps QM/MM relaxation
- PMFs at the DFTB3/ff14SB level

