## Supplementary Information for "Influence of Wobbling Tryptophan and Mutations on PET Degradation Explored by QM/MM Free Energy Calculations"

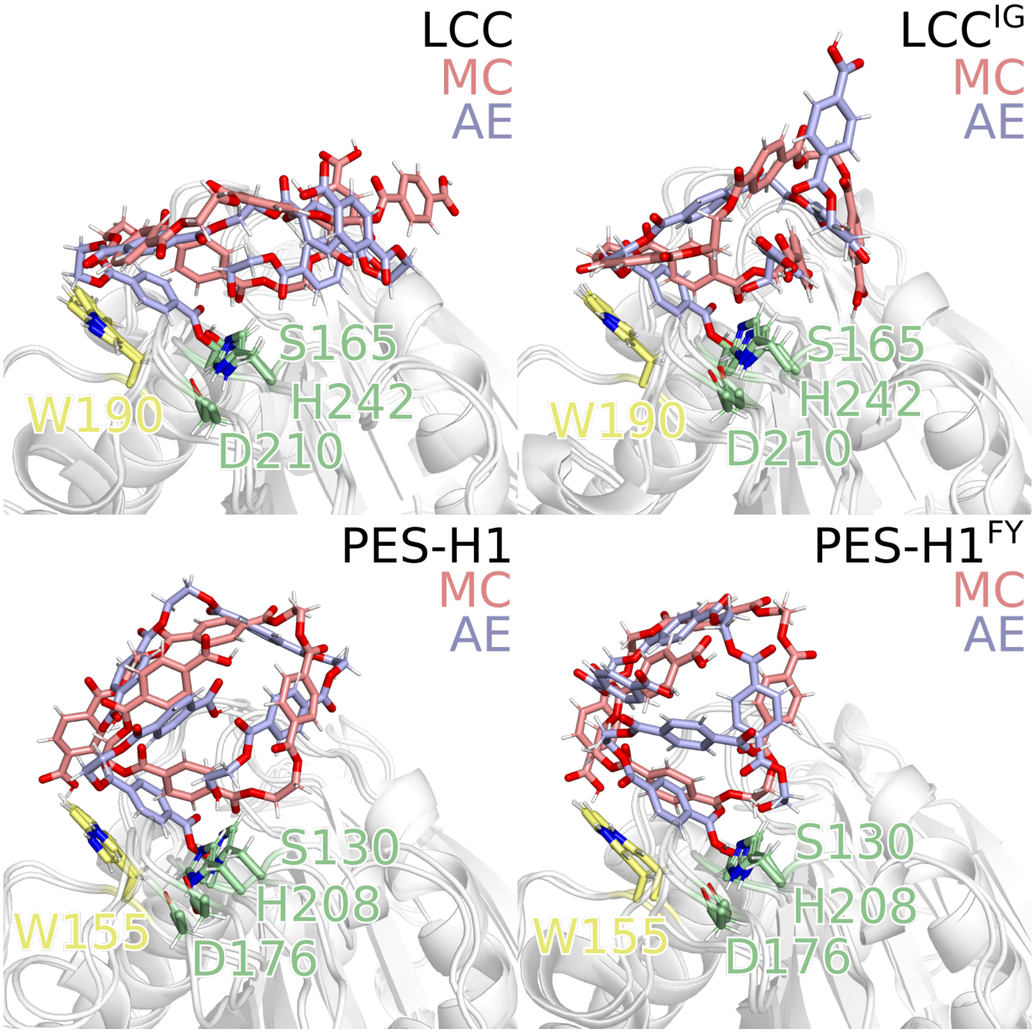


Figure S1: PET conformations of the MC (red) and AE (blue) after 20 ns MM production run for LCC, $LCC^{IG}$, PES-H1 and PES-$H1^{FY}$.

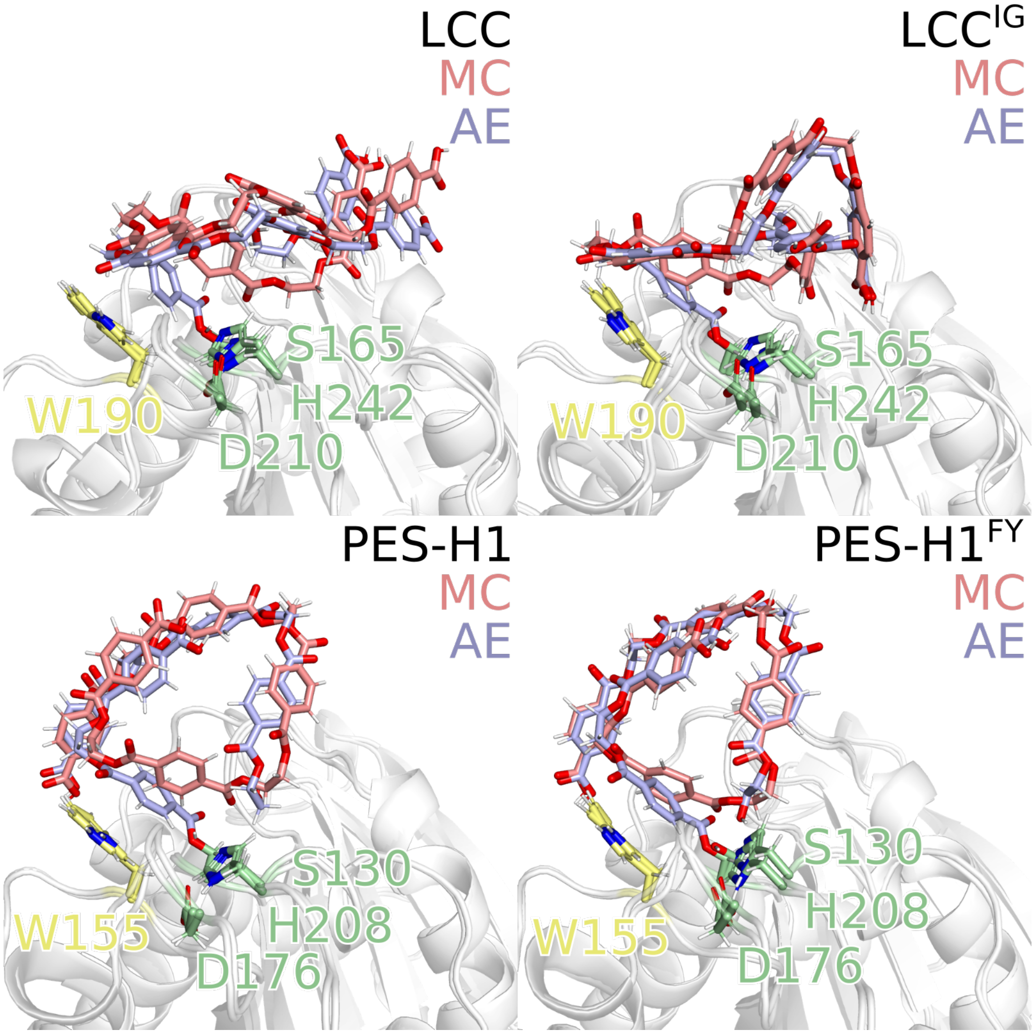


Figure S2: PET conformations of the MC (red) and AE (blue) after 50 ps QM/MM relaxation before conducting ASM for LCC, $LCC^{IG}$, PES-H1 and $PES\text{-}H1^{FY}$.

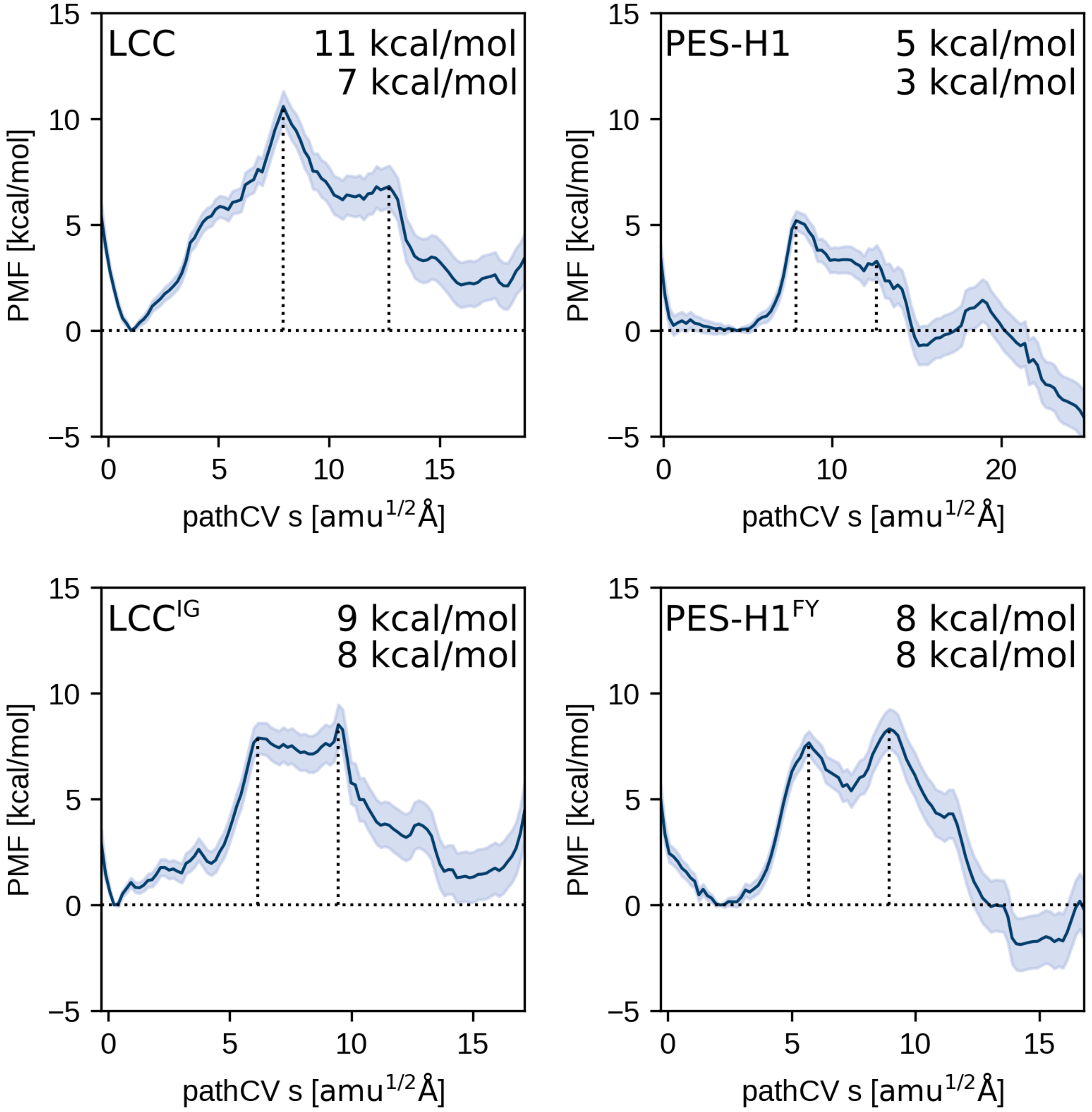


Figure S3: PMF at the DFTB3/ff14SB level of the PET deacylation of LCC (top, left) and PES-H1 (bottom, right) and their highly active variants $LCC^{IG}$ (top, right) and PES-$H1^{FY}$ (bottom, right). The height of the two barriers is provided as well.
